# The Genetic Insulator RiboJ Increases Expression of Insulated Genes

**DOI:** 10.1101/317875

**Authors:** Kalen P Clifton, Ethan M Jones, Sudip Paudel, John P Marken, Callan E Monette, Andrew D Halleran, Lidia Epp, Margaret S Saha

## Abstract

The self-cleaving ribozyme RiboJ is an insulator commonly used in genetic circuits to prevent unexpected interactions between neighboring parts. These interactions can compromise the modularity of the circuit, impeding the implementation of predictable genetic constructs. Despite its utility as an insulator, a quantitative assessment of the effect of RiboJ on the properties of downstream genetic parts is lacking. Here, we characterized the impact of insulation with RiboJ on expression of a reporter gene driven by a promoter from a library of 24 frequently employed constitutive promoters. We show that depending on the strength of the promoters, insulation with RiboJ increased protein abundance between twofold and tenfold and increased transcript abundance by an average of twofold. This result is the first to demonstrate that genetic insulators can impact the expression of downstream genes, potentially hindering the design of predictable genetic circuits and constructs.

## Main Text

A fundamental goal of synthetic biology is the prediction of the behavior of genetic constructs based on the properties of their constituent genetic parts [1]. This predictability relies on the use of modular parts, whose behaviors are unaffected by other parts in the construct [2]. However, modularity can be compromised when interactions between neighboring parts create unintended functional sequences, altering genetic context [3]. These alterations lead constructs to behave unpredictably, impeding their function.

One significant source of these alterations of genetic context is the use of synthetic promoters containing regulatory sequences downstream of the transcriptional start site [4]. This additional sequence is transcribed, leading to the inclusion of unintended nucleotides termed “RNA leaders” at the 5’ end of the transcript. It has been shown that these RNA leaders can modify the stability and secondary structure of mRNA, which alters the translational properties of genetic constructs [2,4]. The nature of these alterations is specified by the interactions between a given RNA leader and the downstream sequence of the transcript. Thus, changes to a construct’s behavior will depend on both the specific promoter used and the composition of a construct.

To circumvent the effects of RNA leaders, constructs can be designed to include genetic insulators, which isolate parts from unwanted interactions with their neighboring regions. One routinely used genetic insulator is the synthetic self-cleaving ribozyme RiboJ [4,5,6,7]. RiboJ is a 75 nucleotide (nt) sequence which is inserted in a construct at the junction of a promoter and its downstream sequence. After transcription, RiboJ self-cleaves, removing the RNA leader and leaving behind a short sequence from its uncleaved region. This elimination of RNA leaders standardizes the behavior of promoters across constructs insulated with RiboJ, aiding the design of predictable genetic constructs.

Although RiboJ is frequently used as an insulator in genetic constructs, there has been limited characterization of what, if any, effect this insulation has on downstream genetic parts. Since the construction of accurate and robust genetic circuits could be impeded if insulation with RiboJ has any unanticipated effects on gene expression, we characterized the impact of RiboJ insulation on the gene expression of a set of constitutive promoter constructs.

We assembled two sets of reporter constructs that differed only by the presence RiboJ. Each set contained 24 constructs, which each expressed a superfolder green fluorescent protein (sfGFP) [4] reporter with a different synthetic *Escherichia coli* (*E. coli)* constitutive promoter part. The collection of promoter parts spans a wide range of transcriptional strengths and includes the well characterized and commonly used Anderson Promoter Library [8], as well as synthetic hybrid promoters such as R0011 (pLlacO-1) **(Supplementary Table 1)**. Each construct was transformed into BL21 *E. coli* and expression of the fluorescent reporter was measured using flow cytometry [9]. Detailed methods can be found in the supporting information.

We found that for each pair of promoter parts, the construct insulated by RiboJ had greater absolute fluorescence than the corresponding construct without RiboJ **(Figure 1a, Supplemental Figure 1)**, with increases ranging from twofold to tenfold **(Figure 1b)**. We found that the fold change in expression with RiboJ exhibited bimodality between “strong” and “weak” promoters. Across 18 of our assay’s 19 strongest promoters, insulation with RiboJ increased absolute fluorescence by an average of sevenfold, ranging from threefold to tenfold. For the remaining promoters, we found that the magnitude of the increase was lower, with an average increase of fivefold, ranging from twofold to eightfold. While we found that RiboJ exhibited bimodality between “stronger” and “weaker” promoters, we did not find a monotonic relationship between promoter strength and fold change in protein expression **(Supplemental Figure 2)**.

**Figure 1:**
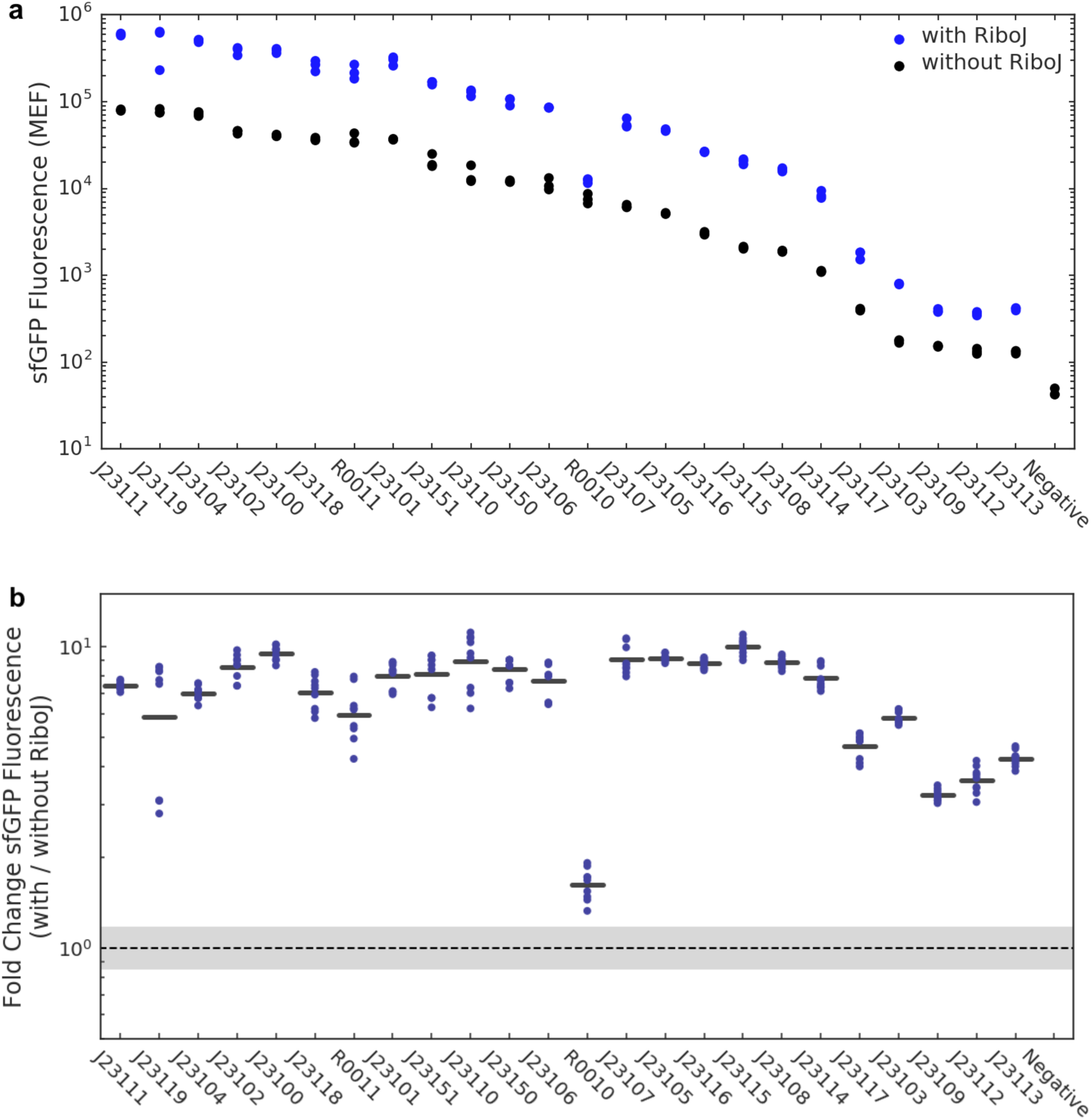
**(a)** Absolute fluorescence of constructs denoted by BioBrick ID (**Supplementary Table 1**), with (blue) and without (black) RiboJ insulation as measured by calibrated flow cytometry. Each dot represents the geometric mean fluorescence of n > 10,000 cells. **(b)** Fold change in fluorescence of constructs when insulated with RiboJ. Bars represent the fold change in the mean fluorescence across replicates, and dots represent all pairwise fold changes between replicates. The dashed line and grey region indicate one geometric SD factor around the geometric mean of a null fold change distribution computed from the fluorescence data (**Supplemental Methods**).

Since the increase in protein expression with RiboJ could be attributed to either differential transcription or translation of insulated genes, we characterized the effect of insulation with RiboJ on the relative abundances of sfGFP transcripts. We used reverse transcription digital droplet qPCR (ddPCR) with Uroporphyrinogen-III C-methyltransferase (CysG) serving as an endogenous reference [9]. We found that insulation of constructs with RiboJ, increased sfGFP transcript abundance by an average of twofold, while there was no change in transcript abundance of our reference gene CysG on average **(Supplemental Figures 3, 4)**. The mean fold change for sfGFP transcript counts was greater than for a null distribution and for an endogenous reference gene, which indicates that the observed increase in the transcript abundance of sfGFP is indeed due to insulation with RiboJ **(Figure 2, Supplemental Figure 5)**.

**Figure 2:**
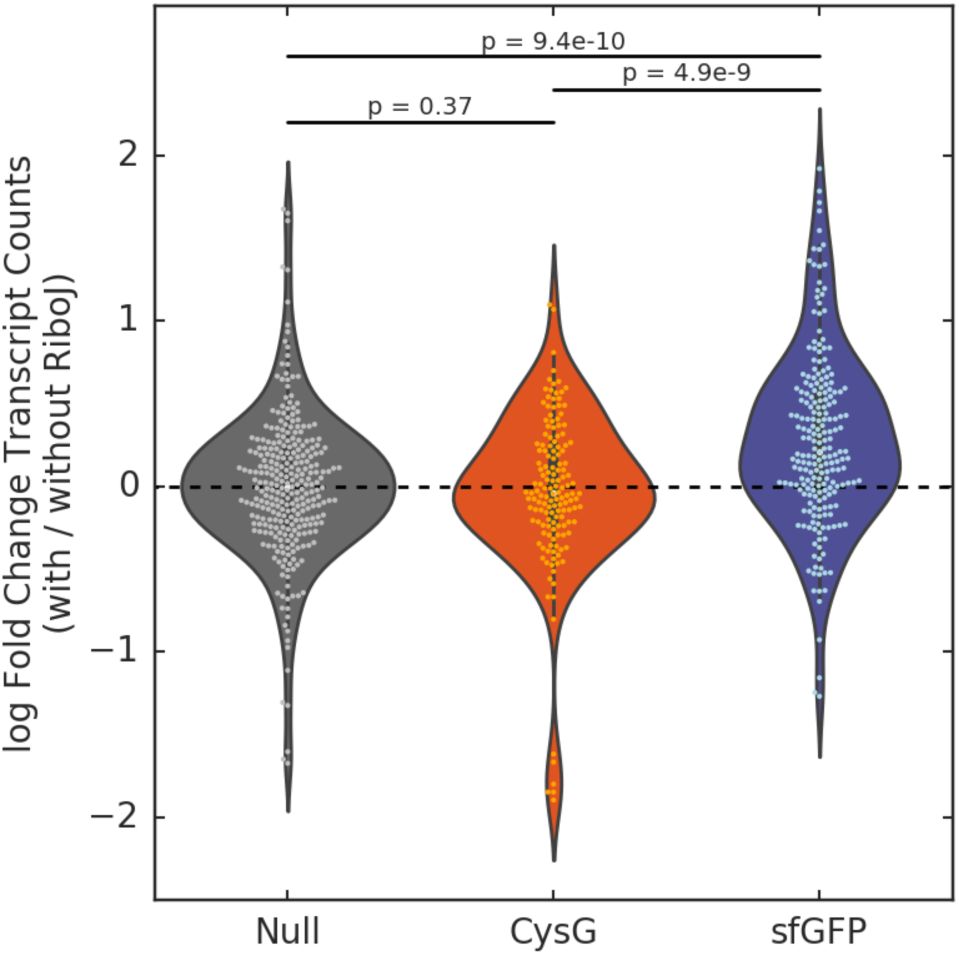
Fold change in the transcript abundance of CysG, sfGFP, and null distribution when promoter constructs are insulated with RiboJ. P values were calculated using Welch’s one-tailed t-tests with hypotheses sfGFP > Null (p = 9.41e-10) and sfGFP > CysG (p = 4.94e-09). For the comparison of Null and CysG, Welch’s two-tailed t-test was used (p = 0.37). Dots represent all pairwise ratios of replicates (9 dots per promoter). The null distribution was calculated by comparing all pairwise ratios of sfGFP transcript counts within a RiboJ condition (with/with or without/without) for a given promoter, excluding identity ratios (where a sample is divided by itself) **(Supplementary Methods)**.

While the cause of the increased expression of RiboJ-insulated genes is unknown, the increased expression levels of protein and mRNA could be driven mainly by an increase in mRNA concentration. This increase in mRNA concentration could be due to increased mRNA stability from the terminal hairpin formed by the remaining RiboJ sequence post-cleavage [4, 11]. We found that sfGFP transcript abundance was well correlated to sfGFP protein concentration **(Supplemental figure 6)**, which could imply that increased protein abundance is driven by a RiboJ-associated increase in mRNA abundance. However, the fold change in transcript abundance for the collection of constructs did not correlate well with fold change in protein **(Supplemental figure 7)**, which suggests that increases in protein abundance due to RiboJ could be due to translational processes as well as increases in transcript abundance. Additional investigation is required to determine the causes of these increases in gene expression under insulated conditions.

The genetic insulator RiboJ is a valuable tool that aids the implementation of predictable genetic circuits by allowing promoter characterization to be standardized across genetic constructs. While insulation with RiboJ is widely used, there has been no comprehensive characterization of its effects on the expression of insulated genes. Here we provide a quantitative characterization of the effect of insulation with RiboJ on a collection of promoter constructs. We determined that insulating a construct with RiboJ leads to an increase in protein expression and transcript abundance. Since unanticipated increases in gene expression can decrease circuit performance by increasing cellular metabolic strain or by causing decreases in the dynamic range of portions of the circuit [12], this characterization provides valuable information for the design and implementation of genetic constructs and circuits.

## Supporting Information

SI.docx: Supporting figures S1-7, Supplemental methods, Supporting Table 1, Construct design

Data.csv: Experimental Data

Analysis.ipynb: Analysis methods

## Acknowledgements

We would like to thank Joseph Maniaci for his assistance in the initial cloning of the construct library. We would also like to thank Vice Provost Dennis Manos for his intellectual support throughout the project.

## Abbreviations

ddPCR: digital droplet quantitative polymerase chain reaction
sfGFP: superfolder GFP

## Author Contributions

ADH designed constructs. EMJ and KPC cloned constructs with help of ADH. EMJ, KPC and SP isolated RNA. LE performed reverse transcription and digital droplet qPCR. EMJ and KPC performed flow cytometry measurements and analysis. EMJ, KPC and JPM analyzed data. KPC and JPM created figures. MSS supervised project. EMJ, KPC wrote manuscript. KPC, ADH EMJ, CEM, JPM, SP, MSS edited manuscript.

